# Stable multi-level social structure is maintained by habitat geometry in a wild bird population

**DOI:** 10.1101/085944

**Authors:** Damien R. Farine, Ben C. Sheldona

## Abstract

Social structure can have profound evolutionary and ecological implications for animal populations. Structure can arise and be maintained via social preferences or be indirectly shaped by habitat structure. Understanding how social structure emerges is important for understanding the potential links between social structure and evolutionary and ecological processes. Here, we study a large community of wild birds fitted with uniquely-coded passive integrated transponder (PIT) tags and recorded on a grid of automated feeders fitted with radio frequency identification (RFID) antennae. We show that both large-scale and fine-scale network communities are consistent across years in this population, despite high generational turn-over. Studying the process that generates community structure, here the movement of individual birds across the woodland, suggests an important role of habitat geometry in shaping population-level social community structure. Our study highlights how relatively simple factors can produce apparent emergent social structure at the population scale, which has widespread implications for understanding eco-evolutionary dynamics.

## INTRODUCTION

The social environment can profoundly shape the life histories of animals. Who individuals associate with can determine the information they have access to (Dall *et al.* 2005; Seppanen *et al.* 2007; Valone 2007), affect how well they can exploit resources (Pruitt & Riechert 2011; Aplin *et al.* 2014; Pruitt & Keiser 2014), and impact their ability to successfully reproduce (Formica *et al.* 2011; Formica *et al.* 2012; Wey *et al.* 2013; Farine & Sheldon 2015). Social structure can also influence population processes, such as the spread of information within (Aplin *et al.* 2012; Allen *et al.* 2013; Aplin *et al.* 2015a) and between (Farine *et al.* 2015a) species, and the spread of disease among individuals (VanderWaal *et al.* 2013; Adelman *et al.* 2015). Finally, individuals can experience selection arising from properties of their social groups (Formica *et al.* 2011; Farine & Sheldon 2015) or their communities (Pruitt & Goodnight 2014). Quantifying the factors that shape individuals’ social environments is a key step in identifying the evolutionary drivers of sociality (Goodale *et al.* 2010; Farine, Montiglio & Spiegel 2015), as the mechanisms that determine where, and with whom, individuals live their lives are what selection arising from population processes (e.g. via disease burden or information use) can act upon.

Social structure (the pattern of connections that emerges from interactions among individuals, often represented using social networks) is generally considered to arise from individuals making decisions about with whom to associate (Hinde 1976; Whitehead 2008; Kurvers *et al.* 2014). The overall structural properties of a population are likely to be emergent outcomes of the combined behaviours and social decisions of all individuals in the population, which for most animals includes heterospecifics. A number of recent studies have highlighted potential multi-level social structure in animal populations (Wittemyer, Douglas-Hamilton & Getz 2005; Archie, Moss & Alberts 2006; Schreier & Swedell 2009), with smaller ‘groups’ or ‘communities’ of individuals being embedded in larger communities containing multiple groups. The implications of such population structure are profound, for example individuals within a community would experience more similar modulating effects of the social environment on selection relative to individuals from different communities (Montiglio, McGlothlin & Farine 2018). However, we still know relatively little about how multi-level societies emerge.

Because spatial proximity is a necessary requirement for associations or interactions (i.e. connections) among individuals in many species of animals (Farine 2015), patterns of connections among individuals are bound to be shaped by a number of extrinsic factors, such as geometry and structure of habitat features that promote or restrict individual movements (Wiens 1976; O’Brien *et al.* 2006; Cox *et al.* 2016; He, Maldonado-Chaparro & Farine 2019). At a local scale, habitat structure, such as the presence of understorey density, can channel individual movements, thus increasing the propensity for contact among individuals. At a broader scale, movement ‘highways’ can significantly reduce the social distance among individuals despite a large spatial separation, and therefore facilitate the flow of information or disease (Brockmann & Helbing 2013). For example, sleepy lizards (Tiliqua rugosa) living in open habitats had fewer contacts with conspecifics than those living in more structured habitats (Leu *et al.* 2016). Whilst habitat features can promote movements, the geometry of habitat (such as the boundary of fragmentized habitat) can also introduce barriers that constrain individual movements. Such barriers can result in a structure where the social distance between two individuals (i.e. their degrees of separation in a social network) on either side of this barrier could be much greater than their actual spatial distance. Habitat geometry and features can therefore impose structure in a social network that could easily be interpreted as arising socially. Given that local heterogeneity in gene flow can lead to rapid evolutionary differentiation (Garant *et al.* 2005), integrating knowledge about fine-scale environmental heterogeneity into studies of social structure could fundamentally alter our understanding of adaptation and the ability for animals to respond to selective pressures.

A major challenge in identifying extrinsic factors that drive population social structure is the need to track the movement of individuals in space, patterns of social connections among the majority of individuals in a population, together with information about the habitat in which they live (Strandburg-Peshkin *et al.* 2017). Recent technological advances have eased the logistical constraints of sampling many individuals moving across large areas (Tomkiewicz *et al.* 2010; Rutz *et al.* 2012; Ryder *et al.* 2012; Krause *et al.* 2013; Finn *et al.* 2014; Kays *et al.* 2015; Levin *et al.* 2015; Strandburg-Peshkin *et al.* 2015; Jacoby & Freeman 2016). In particular, passive integrated transponder (PIT) tags are cheap electromagnetic tags that can be fitted to many individuals at once (Bonter & Bridge 2011), thus overcoming the challenges associated with studying entire animal communities. Because PIT tags do not rely on battery power to emit a signal, but instead are detected by affecting the magnetic field in radio frequency identification (RFID) antennas, they provide the capability to track individuals across years and life stages. Further, advances in the analytical tools, such as social network analysis (Farine & Whitehead 2015) and community detection algorithms (Fortunato 2010), are facilitating greater insight into patterns of social structure. These technological advances have underpinned a boom in the study of animals’ social networks (Krause *et al.* 2015), and in networks formed by the movement of individuals across space (Finn *et al.* 2014; Jacoby & Freeman 2016). However, much less effort has focused on identifying the key mechanisms that shape overall social structure at the scale of populations.

In this paper, we investigate whether habitat structure affects patterns of movement, and in turn drive social structure, in a large population comprising 5 species of wild songbird. A number of previous studies in our population have investigated how genetic traits (Radersma *et al.* 2017), phenotypic traits such as age, personality, sex, and immigration status (Quinn *et al.* 2011; Aplin *et al.* 2013; Aplin *et al.* 2014; Farine 2014; Aplin *et al.* 2015b; Farine *et al.* 2015b), species identity (Farine, Garroway & Sheldon 2012; Farine *et al.* 2014), and diel cycles (Farine & Lang 2013; Hilleman *et al.* in press) affect individual-level movement and social behaviour, and that who individuals associate with can impact their fitness (Farine & Sheldon 2015). Using data from this population collected over 4 consecutive winters, we first quantify overall patterns of movements across the landscape each winter to determine whether individuals move evenly through space, or if there are consistent movement corridors. Based on studies showing that birds are sensitive to open habitat (Quinn *et al.* 2012) (which heighten the risk of predation), we predict that birds should show a preference for moving along corridors comprising denser understory habitat. Second, we use patterns of associations among individuals in each winter to construct social networks, and use these social networks to determine whether social structure is consistent across winters. Given that approximately 50% of individuals in the population each winter are first-year birds, and approximately 50% of first-year birds are immigrants into the population (born outside the study area), we predict that there would be no consistent social structure at local scales. However, because the study area remained identical across winters, we predict that any effects of the habitat on the social structure of the population would be repeated each winter.

## METHODS

### Study location, study species, and population dynamics

The study was undertaken in Wytham Woods, Oxfordshire, UK (51° 46’ N, 01° 20’ W), a 385ha area of broadleaf deciduous woodland ‘island’ surrounded by intensive agriculture. Pairs of birds hold territories during the breeding season (April – June), but form loose fission-fusion groups during the winter, flocking with unrelated individuals that forage for ephemeral food sources. Flocks often contain multiple species (Farine, Garroway & Sheldon 2012), and our study also includes data on the five most common flocking species: great tits (Parus major), blue tits (Cyanistes caeruleus), marsh tits (Poecile palustris), coal tits (Periparus ater), and nuthatches (Sitta europaea). Tits are generally short-lived—great tits have a mean life span of 1.9 years. This short generation time results in high annual population turn-over and inter-annual variation in population sizes. Good breeding conditions lead to large population sizes, whereas poor breeding conditions result in fewer juveniles and a much reduced population size.

### PIT-tagging birds

All birds in the study were caught in either a nest box (as parents and as chicks) or a mist-net (approximately half the population are birds that immigrate). Each bird was fitted with uniquely numbered British Trust for Ornithology metal leg ring, and a uniquely-coded passive integrated transponder (PIT) tag (IB Technologies, UK) that was fully enclosed in a moulded plastic ring fitted to the other leg. Each PIT tag contains a unique code that can be recorded by antennae (see next section), and these were then matched to the bird’s ring number. We ceased fitting PIT tags to coal tits from October 2012 as the tags were aggravating pox lesions on birds legs during a naturally-occurring epidemic. For each bird that was caught and tagged, we recorded the age and sex (where possible following Svensson 1992).

### Data collection

We placed 65 automated feeding stations in a evenly-spaced grid covering the entirety of Wytham Woods and small isolated patches of woodland nearby. Commercial bird feeders (Jacobi Jayne, UK) were fitted with a radio frequency identification (RFID) antenna on each of the lower two access holes and other access holes were blocked. The antennae recorded the unique PIT tag code, time and date for each visit by a marked bird. For the duration of the study, the feeding stations were scheduled to open and begin logging at 6am on Saturday mornings, and shut after dusk on Sunday evening. Feeders were in place December to February in the winter 2011-12 (13 weeks), from December to early March of winters 2012-13 and 2013-14 (14 weeks each), and for January and February in the winter of 2014-15 (8 weeks). This data collection resulted in 49 unique weekends and 98 complete data logging days over the 4 winters. In later winters, 6 feeders covering two external sites were replaced for a separate experiment, and thus the data were not included in these analyses.

### Constructing movement networks

From the logging data, we recorded every case of movement by a bird from one feeding station to another within the same day (a total of 83071 movements over 4 winters), and used these to create movement networks that quantify the connectedness between each pair of feeding stations in the study. We used daily sequential detections to maximize our chances of correctly inferring direct movement pathway (e.g. moving between locations A and D via locations B and C) and minimizing our chances of incorrectly inferring movement pathways (e.g. estimating only a direct movement between A and D). Because the number of movements are inherently linked to the number of individuals present at a specific pair of feeders, we also created a network describing the rate of movement between feeders, where the rate was defined as the probability that an individual at one focal feeder would be observed moving to the other focal feeder within a day.

### Understory habitat density

We used data from Kirby *et al.* (Kirby *et al.* 2014) to quantify the habitat structure between each feeding site. In that study, the authors recorded, among other measures, the shrub cover density (0.5 m to 2.5 m above ground) along the diagonal of 164 different 10 m × 10 m quadrats equally spaced throughout Wytham Woods. Here we use data from the 2012 census, which falls roughly in the middle of our study period, but note that there were no major changes to the habitat across years. To extrapolate from the 164 sites, we generated a surface plot where we extrapolated the data to a 10 × 10 m grid of points using the vgm function in the R package gstat (Gräler, Pebesma & Heuvelink 2016) with a spherical model, omitting fitting the nugget component. The resulting figure accurately captures variation in habitat density based on our knowledge of the study site. To calculate habitat density between each pair of feeding sites, we calculated the mean habitat density (as estimated by the surface) along a 20m-wide transect connecting the two sites.

### Inferring flocks and flock membership

The data logged from the PIT tag detections produces bursts of detections in the temporal data stream. These vary in length depending on the size of the flocks present (which increases during the course of the day, Farine & Lang 2013; Hilleman *et al.* in press). We used a recently-developed statistical tool for analyzing such data involving Gaussian Mixture Models (Psorakis *et al.* 2012) to extract the start and end times for each distinct flock. This machine-learning method statistically fits Gaussian curves of varying sizes to each burst in the data and allocates each record to the distribution, or ‘gathering event’, into which it falls. We have shown that the Gaussian Mixture Model approach provides a more robust estimation of the social network structure than alternative methods (Psorakis *et al.* 2015).

### Constructing social networks

We defined edges in the social network using the simple ratio index (Hoppitt & Farine 2018): 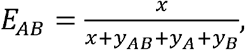, where *E_AB_* is the edge weight between individuals *A* and *B*, *x* is the number of times they were detected in the same flock, *y_AB_* is the number of occasions they were both detected at the same time but not in the same flock, *y_AB_* is the number of detections of *A* where *B* was not seen, and *y_AB_* is the number of detections of *B* where *A* was not seen. The networks for each winter were constructed using *R* package asnipe (Farine 2013).

### Detecting consistent social structure across winters

For each winter’s social network, inferred structural communities in each social network using the edge betweenness community detection algorithm in the R package *igraph* (Csardi & Nepusz 2006). We specified the algorithm to detect *k* = 2 to 65 communities, and recorded the identities of individuals in each community. If communities are structured exclusively by extrinsic factors, we expected a drop in the stability of co-membership by individuals as we created finer grained communities. For example, if a population is spread across three isolated patches of woodland, and birds do not move between woodlands, then we expect that birds will always occur within the same three communities (one for each woodland patch) each winter. By contrast, if communities are structured socially, then we expected smaller communities (local cliques) to be more stable, but little stability in global structure. For example, for a territorial pair-living species living in a lattice-like uniform environment, an algorithm will be able to isolate each pair when identifying *k* = N/2 communities, whereas the communities detected for smaller *k* values will be essentially random.

### Linking movements to community structure

To test for a link between community structure and movement networks, we allocated individuals to their most common feeder, and for each of the feeding stations, selected the community of which the majority of individuals present were members. This enabled us to create a community label for each feeding station (and for each value of *k*), and link these to the network formed by the movement of individuals (Figure 1). To test whether the movement network shaped the community structure in the network for a given value of *k*, we quantified the assortativity coefficient of the network using communities as discrete trait values (Farine 2014; Shizuka & Farine 2016). Assortativity is the measure of how well connected alike nodes are compared to how well connected dislike nodes are, ranging from 1 (all edges connect nodes with the same traits) to −1 (all edges connect nodes with different traits).

**Figure 1:**
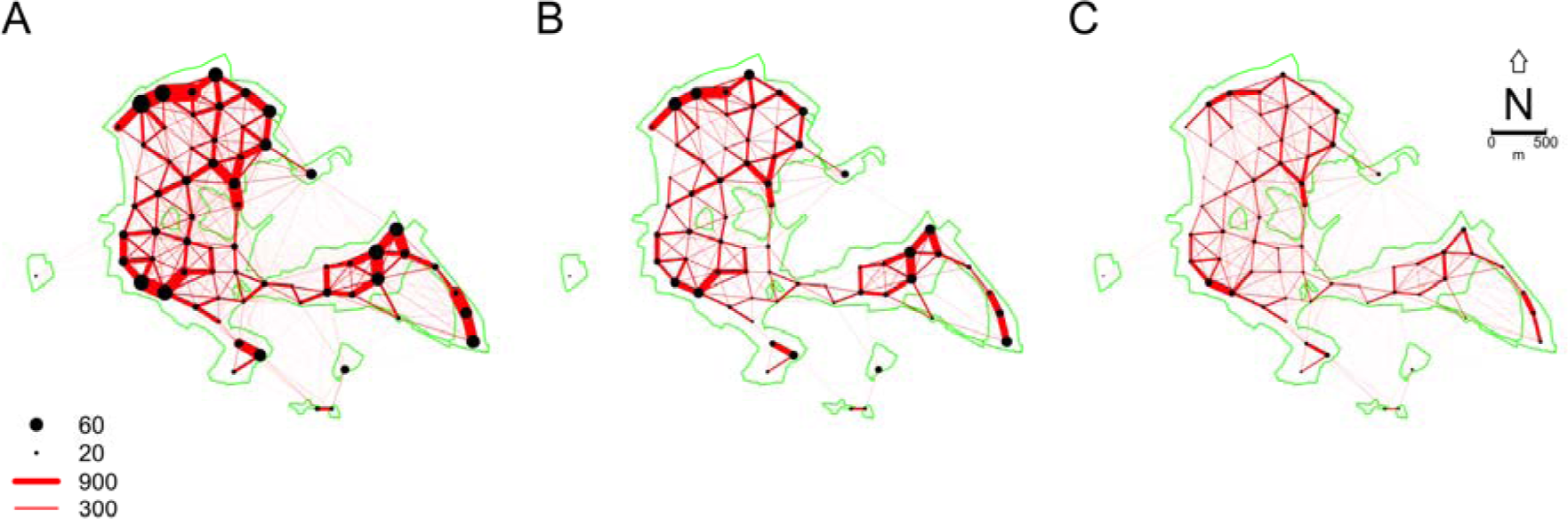

Total movements of (A) all birds, (B) adults, and (C) juveniles (first winters) from all species over 4 winters of data. The thickness of each line represents the number of observations of a bird detected at two feeding stations (black points) in the same day. The size of the points represents the average number of individuals observed at each feeding station. The green outline represents the outline of the wooded areas, which are surrounded by open agricultural land. Note that there is very little movement of birds between Wytham Woods and the four external woodlots, highlighting how closed this population is during the winter. See Figure S1 for statistical evidence that juveniles make significantly more long-distance movements.

## RESULTS

### Movement networks

We first quantified the daily movement rates between different feeding stations. Our population consisted of a total of 6299 unique PIT-tagged individuals, of which 2230 were great tits, 3304 were blue tits, 237 were marsh tits, 169 were coal tits, and 78 were nuthatches. We detected 83701 movements, detections at two feeders on the same day, across all winters (Figure 1). Overall, birds moved very little—our analysis was based on 9612 ‘bird winters’, meaning that each bird moved (on average) fewer than 10 times over a whole winter. We found that most movements occurred between feeding stations where more birds were present.

Not all birds followed the same pattern. Birds in their first winter moved more often than older birds: first-year birds accounted for 61% of all movements, despite making up only made up 38% of the population on average. Of the juveniles that moved, each made on average 20.8 moves, which was significantly more than the average 12.9 moves made by adults detected moving at least once (GLM: β±SE = 0.477 ± 0.007, z = 67.08, P < 0.001, family = Poisson). Further, their movement networks included more long-distance movements than those of adults, resulting in a significantly higher average movement distance (β±SE = 29.5 ± 1.35, t = 21.77,P < 0.001, see also Figures 1 and S1).

We also found species-level differences. Great and blue tits, which comprise the large majority of individuals in the population, exhibited similar movement patterns (difference = 0.5 m, P = 0.520, see Figures S2-3). By contrast, marsh tit movements were typically much more localized (difference blue tit – marsh tit: 48.8 m, P < 0.001; difference great tit – marsh tit: 29.3 m, P < 0.001; see Figures S2-3). Overall, age had the greatest effect on the distribution of movements, with some differences among species.

### Do understory density or habitat geometry predict movement propensity?

We compared the patterns of movement observed across winters, and related these to spatial distance and habitat features. We first re-defined edges in the movement network as the probability that an individual detected at either site moves between them on a given day. Edge weights ranged from 0 representing no movement between adjacent sites to 0.83 representing that each individual detected had an 83% chance of moving between the two sites on a given day (see Figure S4). Weighted multiple regression quadratic assignment procedure (MRQAP)(Dekker, Krackhardt & Snijders 2007) revealed that the movements by birds between feeding stations were significantly more similar from winter-to-winter than expected by chance, even when accounting for distance through the forest and habitat structure (see Table 1, Figure 2). In all winters, birds were significantly more likely to move between ‘close’ feeding stations than distant ones. However, contrary to our predictions, we found no effect of understory habitat density on the propensity for birds to move between feeders.

**Figure 2:**
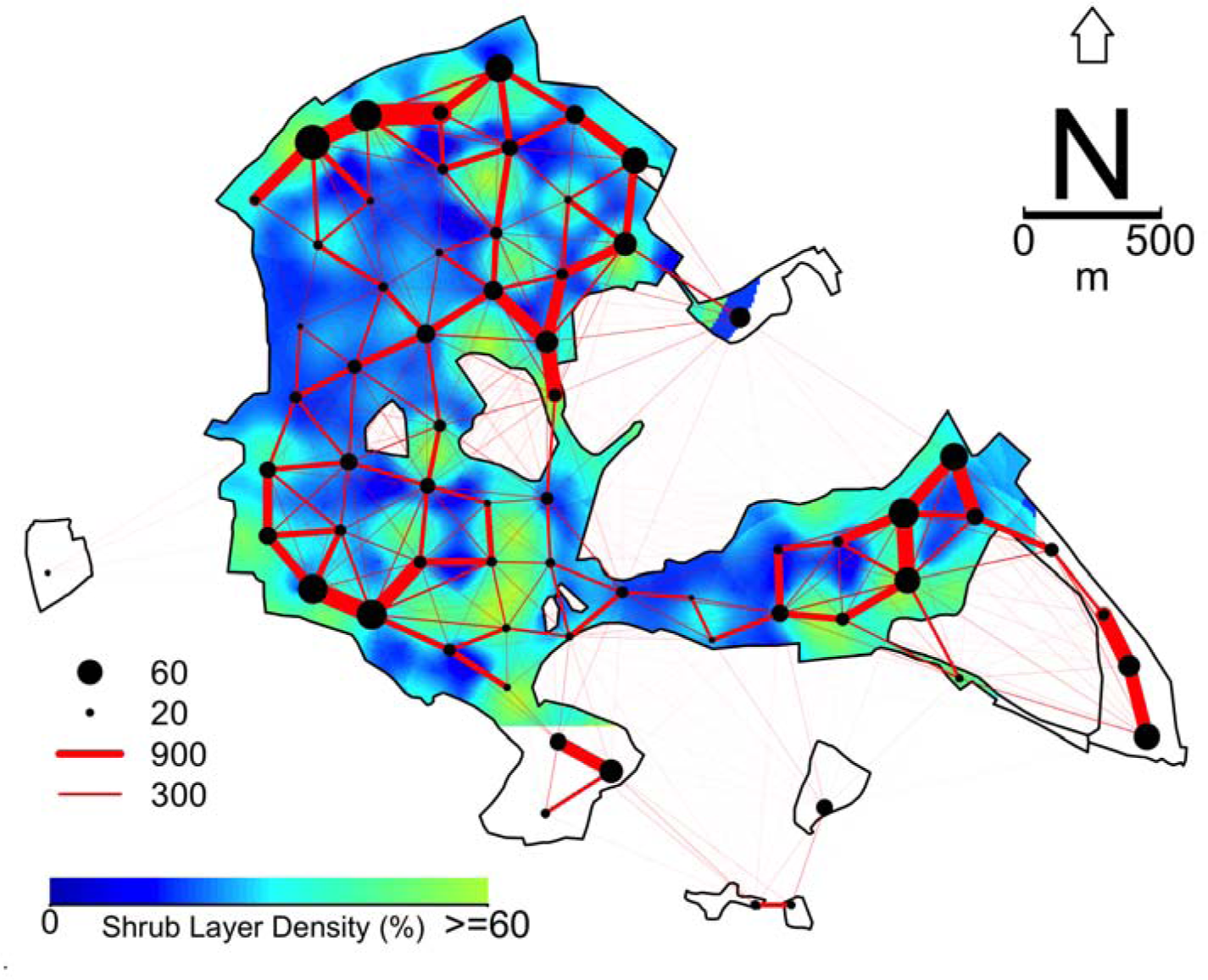
Total movement of all birds (from Figure 1A) overlaid on the understorey habitat density (0.5 m to 2.5 m above ground) calculated at 164 equally-spaced quadrats (see Methods). Areas within the polygon with a white background have no habitat data available, and any links that included such areas were excluded from the analyses. Areas outside of the black polygons are open agricultural land. Blue areas have a largely open understory, whereas green areas have very thick understory. The vast majority of Wytham Woods is closed canopy.

**Table 1:** Results of multiple regression quadratic assignment procedure used to test whether previously observed patterns of movement (probability of moving between sites per capita), distance between sites (through the forest), and habitat density (percentage cover) between sites explain the observed patterns of movement. Bold values represent significant coefficients, * represents significance at P < 0.01, ** represents significance at P < 0.001, based on a two-tailed test. All variables are scaled to 0 mean and unit variance to enable comparison between effect sizes.

| Winter | Previous Winter | Forest Distance | Habitat Density |
| --- | --- | --- | --- |
| 2013 | **0.903\*\*** | **-0.028\*** | -0.010 |
| 2014 | **0.877\*\*** | **-0.060\*\*** | -0.010 |
| 2015 | **0.787\*\*** | **-0.091\*\*** | -0.019 |

Because for many pairs of feeding stations the geometric distance and forest distance are very similar see Figure 2), we used this information to also test whether birds were less likely to move between feeding sites that were separated by non-forest. We did this by calculating the difference between forest distance and Euclidian distance. We found that birds moved less between feeders with a larger difference in distance (i.e. the path through the forest was much longer than the straight-line path), suggesting that birds are avoiding crossing open habitats (Table 2). However, the propensity to move between feeders observed in previous winters was consistently the strongest predictor of future movements. The coefficient values for movements predicted by the previous winter were typically an order of magnitude larger than those of other predictor variables, suggesting that additional undetected factors—potentially social—are driving patterns of movements by birds across this woodland.

**Table 2:** Results of multiple regression quadratic assignment procedure used to test whether birds moved less between feeding sites separated by open space. For each pair of sites, we calculated the difference between the forest distance and the Euclidian distance. Bold values represent significant coefficients, * represents significance at P < 0.01, ** represents significance at P < 0.001, based on a two-tailed test. All variables are scaled to 0 mean and unit variance to enable comparison between effect sizes.

| Winter | Previous Winter | Relative Distance |
| --- | --- | --- |
| 2013 | **0.919\*\*** | -0.001 |
| 2014 | **0.906\*\*** | **-0.019\*** |
| 2015 | **0.828\*\*** | **-0.029\*** |

### Does the population have consistent social structure?

Population-level social structure can have significant implications for population processes. For example, high levels of clustering can reduce the spread of disease within populations (Eames 2008). Thus far, we have shown that habitat geometry, as well as additional unknown factors, contribute to consistent patterns of animal movements. We next investigated how these patterns contribute to the emergent structure of the mixed-species population.

We uncovered two scales that maximized the propensity for pairs of individuals observed in the same community in winter *t* to be observed in the same community in winter *t* + 1. When social networks were partitioned into 2 or 3 communities (Figure 3), individuals observed in successive winters were seen in the same community each winter approximately 90% of the time. These macro communities largely reflect the geometry of the study site, with two core habitats (north-west and the east), and a smaller patch of forest to the south, that is only attached by a narrow neck of vegetation, representing a third community (Figure 4). Specifying the algorithm to detect 4 communities significantly decreased the probability of individuals being re-observed in the same community (no overlap in the ranges in Figure 3), suggesting that there is no stable 4^th^ community. Partitioning the network further by specifying the algorithm to detect more than 4 communities then increased the probability that two individuals observed in successive winters were observed in the same community. Thus, the patterns of social organization at both the population scale (2-3 communities) and at a local scale (>30 communities) were the most stable winter-to-winter. This result suggest that multiple levels of community structure exist in this population, with micro communities nested within macro communities.

**Figure 3:**
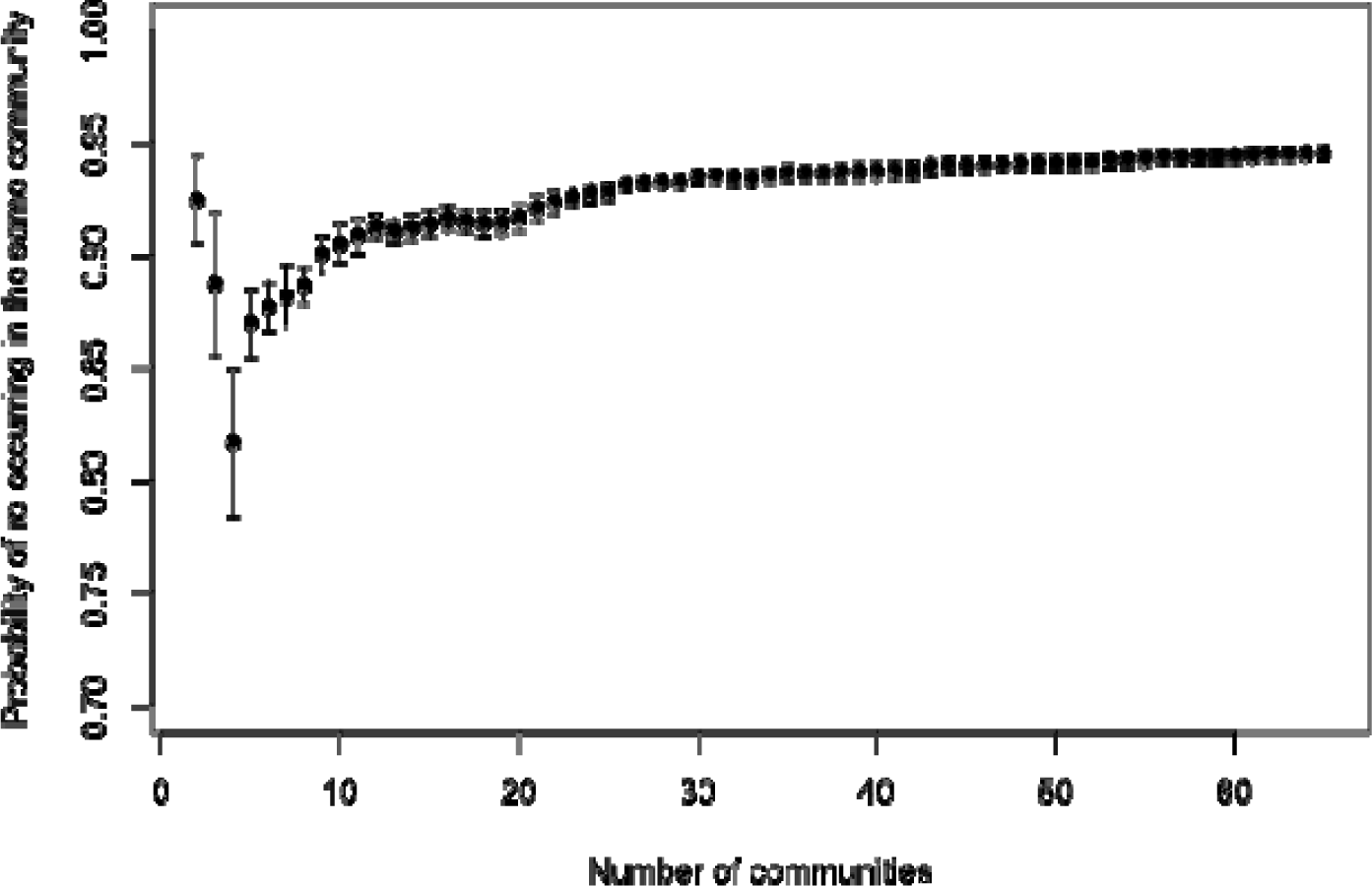

The probability that two birds observed in the same community in one winter remain in the same community in the following winter, given that both are observed. Points represent the mean, and lines represent the range from the 3 pairs of winters. The probability is calculated with the social network partitioned into the same specified number of communities (2 to 65) in all winters.

**Figure 4:**
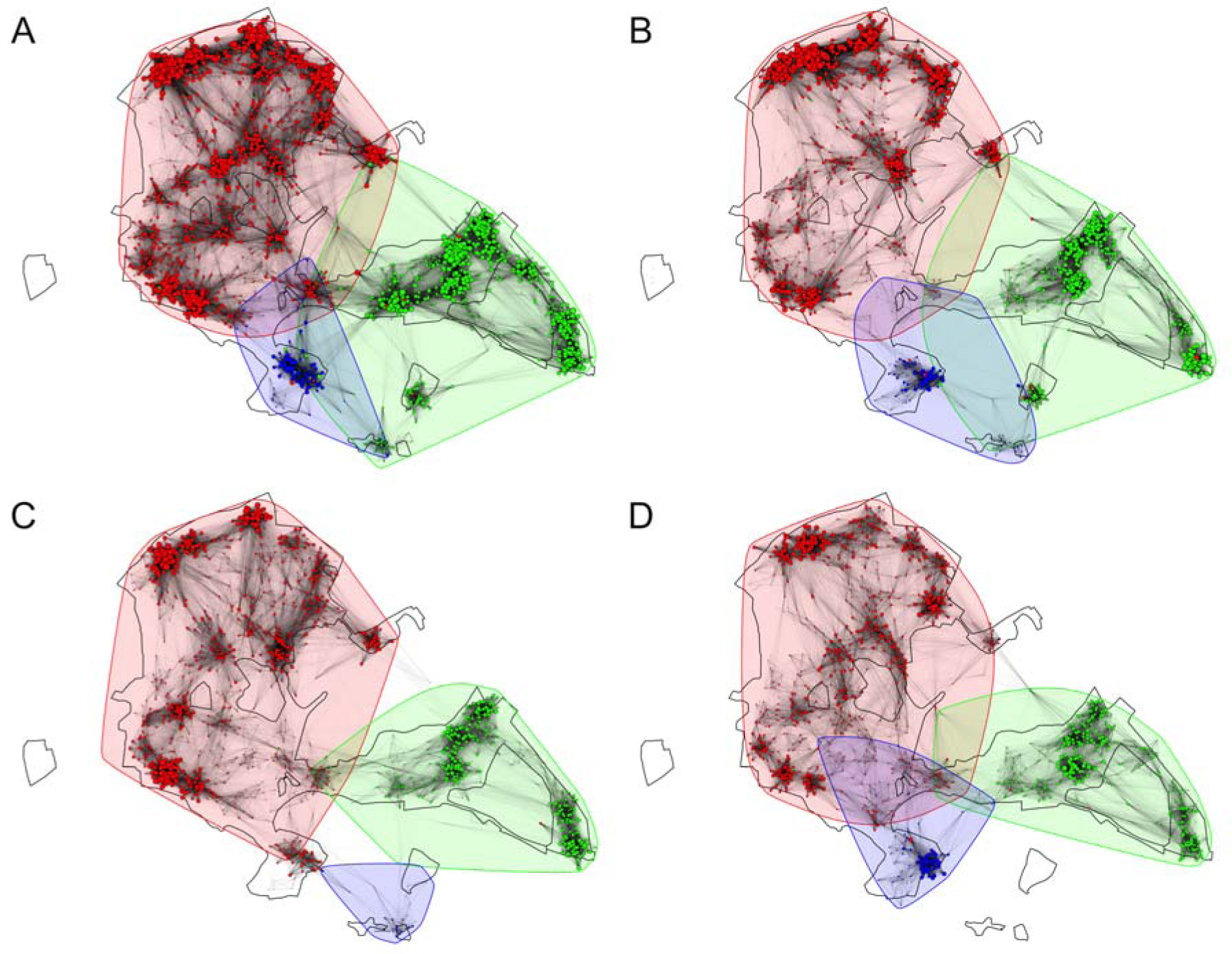
The social network for each winter of the study when partitioned into 3 communities. Each point represents one individual (N = 3019, 2598, 2294, 1701 respectively), and the colours represents the community each node is assigned to. The size of each point represents its weighted degree (larger points have more and/or stronger connections to other individuals). Points are drawn at the average location that the individual was observed, with a small amount of jittering added to reduce the overlap between individuals observed in the same location. Community memberships were inferred using the edge betweenness algorithm applied to each winter independently with the number of communities set to 3.

Partitioning the network into a larger number of communities did not result in one community per feeding station, instead several large communities were maintained and many small, spatially-overlapping, ones were created (Figure 5). Finally, we found no evidence that the composition of species in communities changed based on how many communities were created (Figure S5). Thus, the partitioning of the network into more (micro) communities did not segregate individuals into species-specific clusters, and so stable community structure at a local scale was not explained by simple species-level processes. We note here that there is extensive evidence of social processes driving community structure at these local scales (Krause *et al.* 2015). Such studies of animal social networks are increasingly accounting for effects such as individual home-ranges using null model (Farine 2017). However, fewer studies have explored links between home ranges and community structure (but see Shizuka *et al.* 2014). Our study highlights the importance of exploring this link.

**Figure 5:**
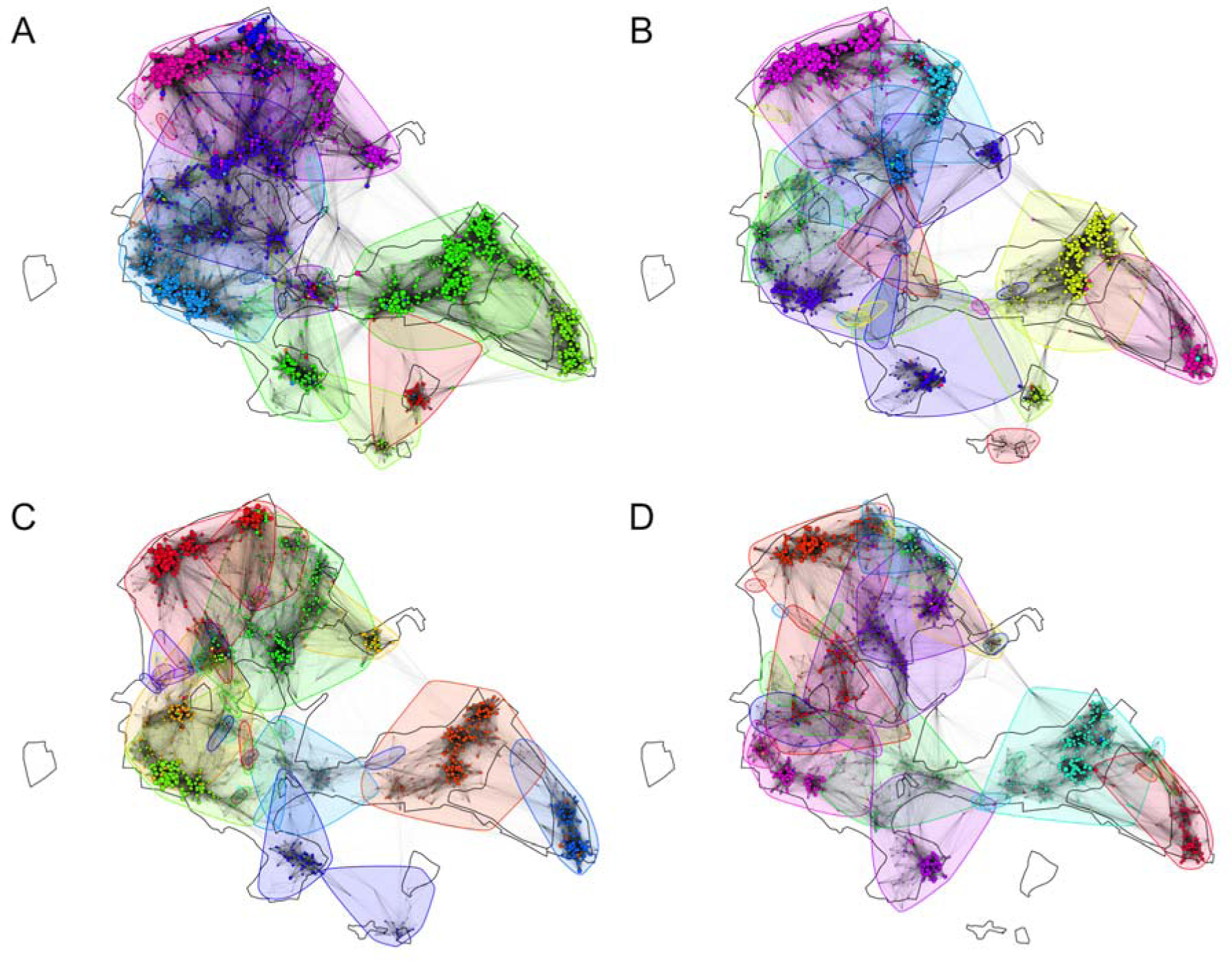
The social network for each winter of the study partitioned into 65 communities. Each point represents one individual, and colours represents the community each node is assigned into (per Figure 4). The size of each point represents its weighted degree (larger points have more and/or stronger connections to other individuals). Points are drawn at the average location that the individual was observed, with a small amount of jittering added to reduce the overlap between individuals observed in the same location. Community memberships were inferred using the edge betweenness algorithm applied to each winter independently with the number of communities set to 65.

### Linking movement patterns to community structure

We investigated whether the regular movements of individuals between particular feeding stations were responsible for global community structure. Individuals living at two locations with frequent movements of individuals between them will be more likely to be connected in the social network, and therefore more likely to be in the same community and share similar social environments. We found that when we partitioned the network into few communities, almost all of the movements were between feeders in the same community (Figure 6). This supports our hypothesis that extrinsic large-scale habitat features shape the broad patterning of the community (i.e. the presence of 2-3 distinct clusters of individuals, see Figure 4) via individual movement (i.e. by disconnecting feeders not directly connected by forest). However, at more local scale, we found that the assortativity coefficient decreased (Figure 6). Thus, as the social network is partitioned into more communities, movements between sites explained less of the community structure, despite the fact that individuals become more likely to re-occur in the same communities across winters (see Figure 3).

**Figure 6:**
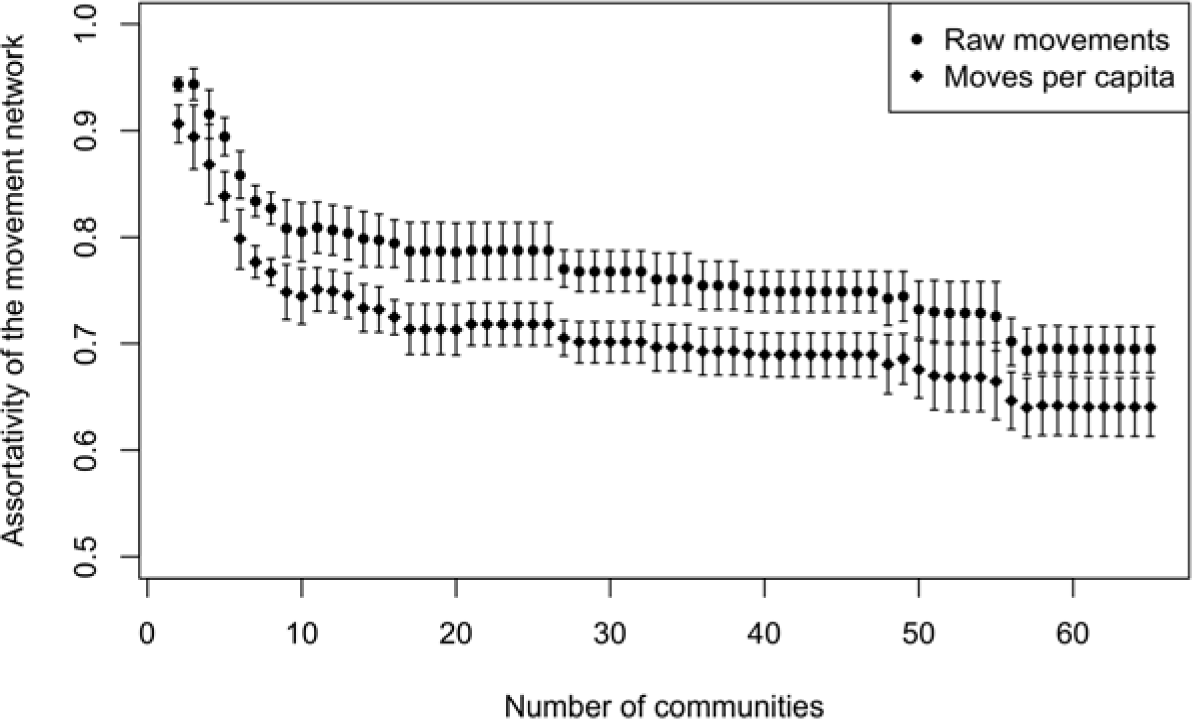
The correlation of movements between feeding stations and community structure decreases with increasing numbers of communities. Each feeding station is allocated to the community in which the majority of individuals are members and used as a trait value to calculate assortment using the raw movement networks (Figure 1A) and the per capita movement network (Figure S2A). High values represent stronger connections between feeding stations in the same community. Points represent the mean, and lines represent the range across the 4 winters.

## DISCUSSION

Our study revealed two levels of social structure, maintained across winters, in a large population of wild birds containing thousands of individuals of five species. At a broad scale, the social network contained two or three communities that were predicted by the regular movement paths used by birds. The movement of birds through the woodland were repeatable each winter, but the similarity in movements across winters was only partly explained by the geometry of the study area. Our analyses suggest, in fact, that some other processes, potentially social processes such as the persistence of local traditions (Mueller *et al.* 2013; Aplin *et al.* 2015a; Jesmer *et al.* 2018), may also be involved. If that is the case, then broad-scale social structure could be, in part, the result of a socially-transmitted inter-generational effect. At a more local scale, we found highly stable social structure, with micro communities of individuals comprising all five species re-associating each winter to maintain consistent communities. Our study thus highlights how factors operating at different scales can shape the social ecology in a wild bird population.

The link between extrinsic habitat factors and community structure in animal populations has been investigated before. For example, community and sub-community structure in Galapagos sealions *Zalophus wollebaeki* are largely driven by the structure of male territories (Wolf *et al.* 2007). However, territorial behaviours are unlikely to play a major role in structuring the winter population of birds in Wytham Woods because the majority of individuals were great tits and blue tits, which are both non-territorial during the winter, although there is some evidence of linkages between summer and winter social structure (Farine & Sheldon 2015; Firth & Sheldon 2016). What is more likely is that individuals have relatively small homeranges that overlap with many other individuals, and that these are not uniformly distributed across the habitat. In non-territorial wintering golden-crowned sparrows (*Zonotrichia atricapilla*), social network communities were also found to be consistent across winters at very small spatial scales (Shizuka *et al.* 2014). The surprising aspect from that study was that golden-crowned sparrows exhibit such stability despite having migrated a long distance from their breeding areas. This aspect is partly replicated in our tit population as approximately 50% of the individuals in any given winter are first-year birds and most birds leave the study area during the summer (Matechou *et al.* 2015). The phenomenon observed in both tits and sparrows suggests that winter sociality is likely to play an important role that goes beyond simple group size effects, and thus could have carry-over effects into the territory structure (Firth & Sheldon 2016) and breeding performance in the following spring (e.g. Farine & Sheldon 2015).

The unexplained consistent structural patterns in both our study and in the golden-crowned sparrow study (Shizuka *et al.* 2014) could represent local traditions that are passed on through social learning. In this scenario, juvenile and immigrant birds copy the movement behaviours of older, resident, birds. This could explain why patterns remained consistent over 4 winters, well beyond the generation time of tits (typically <2 years). Such findings would not be unprecedented. For example, one study in whooping cranes found that juveniles socially learn the migration routes from older individuals (Mueller *et al.* 2013). A previous study in our population showed that social learning can easily lead to persistent local cultures, and that the presence of experienced individuals facilitates the rapid adoption of new behaviours by the next generation (Aplin *et al.* 2015a). The role of such traditions in shaping animal movements and subsequent community structure warrant much further investigation.

Another potentially important feature that we extracted in our study was differential movement patterns between classes of individuals. We found that juveniles typically made more long-distance movements than adults (Figures 1 & S1). This pattern, which is likely to be linked to juvenile dispersal behaviour, has a number of implications for social processes. To overcome strong seasonal changes in the environment, juvenile tits rely on learning from adults in their local environment (Slagsvold & Wiebe 2007). These juveniles therefore play an important role in shaping the overall structure of the social network, and could play a major role as transmission vectors. By coming into contact with a greater number of individuals, they could facilitate the spread diseases or pathogens across communities (as suggested in humans, Del Valle *et al.* 2007), or even introduce novel behaviours into populations.

By investigating the stability of community structure at different scales, we found evidence that tits in Wytham Woods live in a multi-level community structure. Multi-level community structure occurs when animals form small groups, or clusters, of individuals with whom they associate most strongly, and larger groups in which these clusters are embedded. There is increasing interest in multi-level community structure as it can have major implications for how social processes occur (Bell & Ford 1986; de Silva & Wittemyer 2012; Grueter *et al.* 2012; Whitehead *et al.* 2012). Multiple factors can shape the movement (or not) of individuals among social units. These factors can be broadly split into two categories: social factors [such as relatedness (Archie, Moss & Alberts 2006; Croft *et al.* 2012; Godfrey *et al.* 2014), cultural similarity (Cantor *et al.* 2015), or species identity (Bell & Ford 1986)] and habitat factors [features of the environment that modulate where in the environment individuals are found (Croft *et al.* 2003), how they move, and thus whom they encounter (He, Maldonado-Chaparro & Farine 2019)]. Although recent studies have begun to tease apart social versus habitat factors that determine the patterns of contact among individuals with different phenotypic characteristics (Farine *et al.* 2015b), little is known what drives the emergence of global population-level structure. Cantor *et al.* (2015) used simulations to suggest that multi-level communities can emerge when individual segregate into clans formed around similar cultural behaviours. In our study, we found evidence that both environmental and social factors contribute to producing a hierarchical community structure. The general geometry of Wytham Woods is likely to have introduced a repeatable set of large-scale communities (Figure 3). Thus, the shape of the forest is plays a major role in how the population is broadly structured (Figure 4). By studying birds that form mixed-species communities, our study highlights that hierarchical community structure can be the by-product of external processes. Further, the majority of individuals in our study were great tits and blue tits. These birds moved in similar ways, and we could not decompose our networks into groups of one versus the other. Thus, while social mechanisms, such as social preference (Farine *et al.* 2015b) and phenotypic drivers (Croft *et al.* 2003; Croft *et al.* 2009; Aplin *et al.* 2013), can play a large role in determining who individuals affiliate with, the woodland geometry and the resulting behaviour of all individuals combined can generate large-scale static population structures.

The presence of multi-level community structure can have implications for evolutionary dynamics of populations. First, restricted movement can reduce gene flow and lead to divergence in the evolutionary trajectories of sub-parts of each population. Garant *et al.* (2005) demonstrated that differential dispersal reinforces local variation in selection for nestling body mass. In their study, they found that trends in phenotypic variance for body mass in nestlings were very different in the eastern sectors of Wytham Woods and the northern sectors. These two areas represent the two largest population-level communities we found in our study. Second, individuals in the same community will have more similar social environments than individuals occurring in different communities. Thus, any social effects arising via the social environment, such as indirect genetic effects (Moore, Brodie & Wolf 1997), could accelerate patterns of divergence within single populations. Finally, the social environment itself can act as an agent of selection (Wolf & Moore 2010; Farine, Montiglio & Spiegel 2015), and therefore processes that shape social structure are likely to impact the overall strength and direction of selection experienced by populations (Montiglio, McGlothlin & Farine 2018).

One potential limitation of our study is that we employed bird feeders to detect the presence of birds. This approach is what enabled us to collect information on so many individuals simultaneously, but in doing so, we could have also influenced the behaviour of birds. There are three reasons why we do not think that the presence of bird feeders impacted our results. First, the feeders were all identical and open at the same time, meaning that there would be little reason for a bird to choose to relocate to another feeder (and as noted, most birds did not move, and birds moved on average only once every two study days). Second, feeders were evenly spaced out on a grid, meaning that, in the absence of habitat or behavioural heterogeneity, movements between any pair of adjacent feeders should be equally likely. Finally, feeders were open only two days per week, and remained shut for 5 consecutive days. This means that birds would have had to maintain their natural foraging behaviour rather than adapting to a new regime (noting that tits can lose up to 10% of their body weight in a single winters’ night, Owen 1954).

Together the findings from our study highlight several ways in which stable social structure can be maintained in populations. The combination of strong clustering together with some random movements in networks can facilitate the spread of disease or information through the network (Eames 2008). This prediction is supported by the rapid spread and establishment of novel traditions (Aplin *et al.* 2015a) in this population. At the same time, consistent population social structure can lead to phenotypic and genotypic divergence (Garant *et al.* 2005), with potential implications for how animals can adapt to changing environmental conditions. Integrating information about animal social structure with data on both short-term and long-term selective events could yield novel insights into the evolution of social behaviour. As our study highlights, determining the capacity for populations to respond to selective pressures will require an understanding of a range of different drivers that could shape their social structure.

## ACKNOWLEDGEMENTS

We thank Keith Kirby for discussion on the habitat data he collected, and for making these data available to us. We also thank the large number of contributors to the data collection, in particular to Keith McMahon.

