## Supplemental materials for "Stable multi-level social structure is maintained by habitat geometry in a wild bird population"

for

Damien R. Farine*^a,b,c,d^, Ben C. Sheldon^a^

^a^ Edward Grey Institute of Field Ornithology, Department of Zoology, University of Oxford, South Parks Road, Oxford OX1 3PS, United Kingdom.

^b^ Department of Collective Behaviour, Max Planck Institute for Ornithology, Universitätsstrasse 10, 78457 Konstanz, Germany.

^c^ Chair of Biodiversity and Collective Behaviour, Department of Biology, University of Konstanz, Universitätsstrasse 10, 78457 Konstanz, Germany.

^d^ Center for the Advanced Study of Collective Behaviour, University of Konstanz, Universitätsstrasse 10, 78457 Konstanz, Germany.

* Corresponding author:

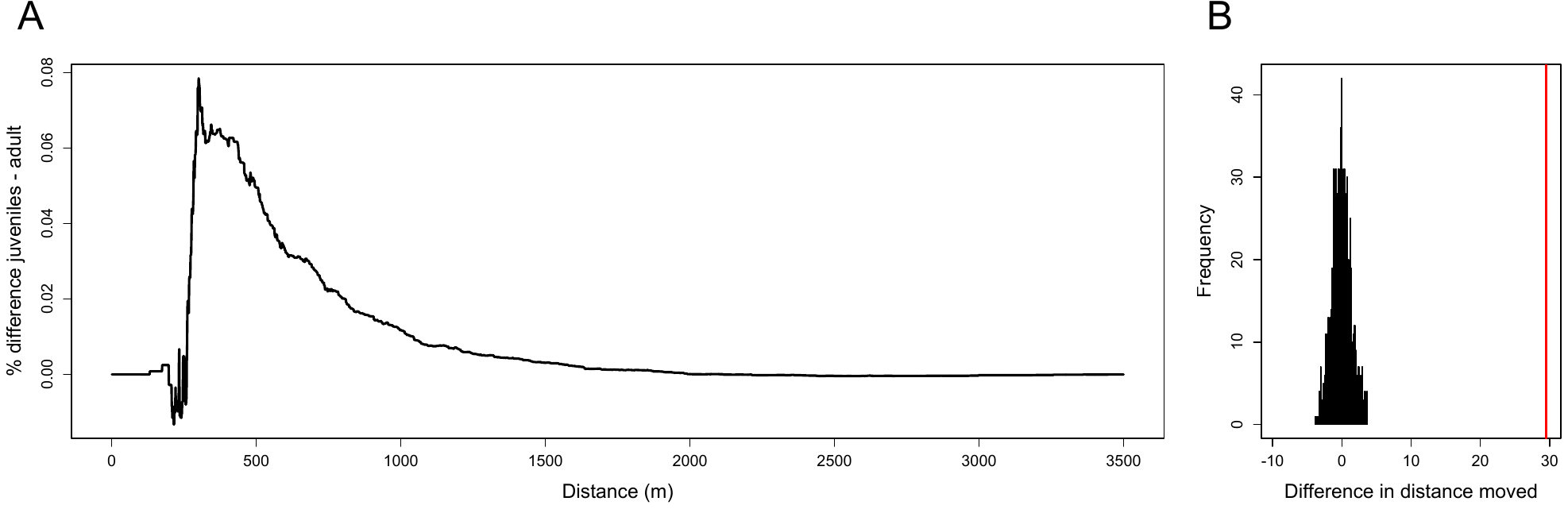

**Figure S1:** Analysis of the difference in movement distance between juveniles and adults. (A) The difference in the percentage of movements for different minimum distance thresholds. Up to approximately 250m (the approximate distance between adjacent feeders), there are disproportionately more movements made by adults. As the minimum distance threshold increases, the difference in the percentage of all data becomes relatively higher for juveniles, meaning that more of the data in juveniles represents large movements. (B) The overall difference in mean distance moved between juveniles and adults (red line) is significantly larger than expected by chance (black histogram), where the null distribution is drawn from 1000 randomisations in which the complete data set was shuffled each time (thus breaking the link between age class and distance moved).

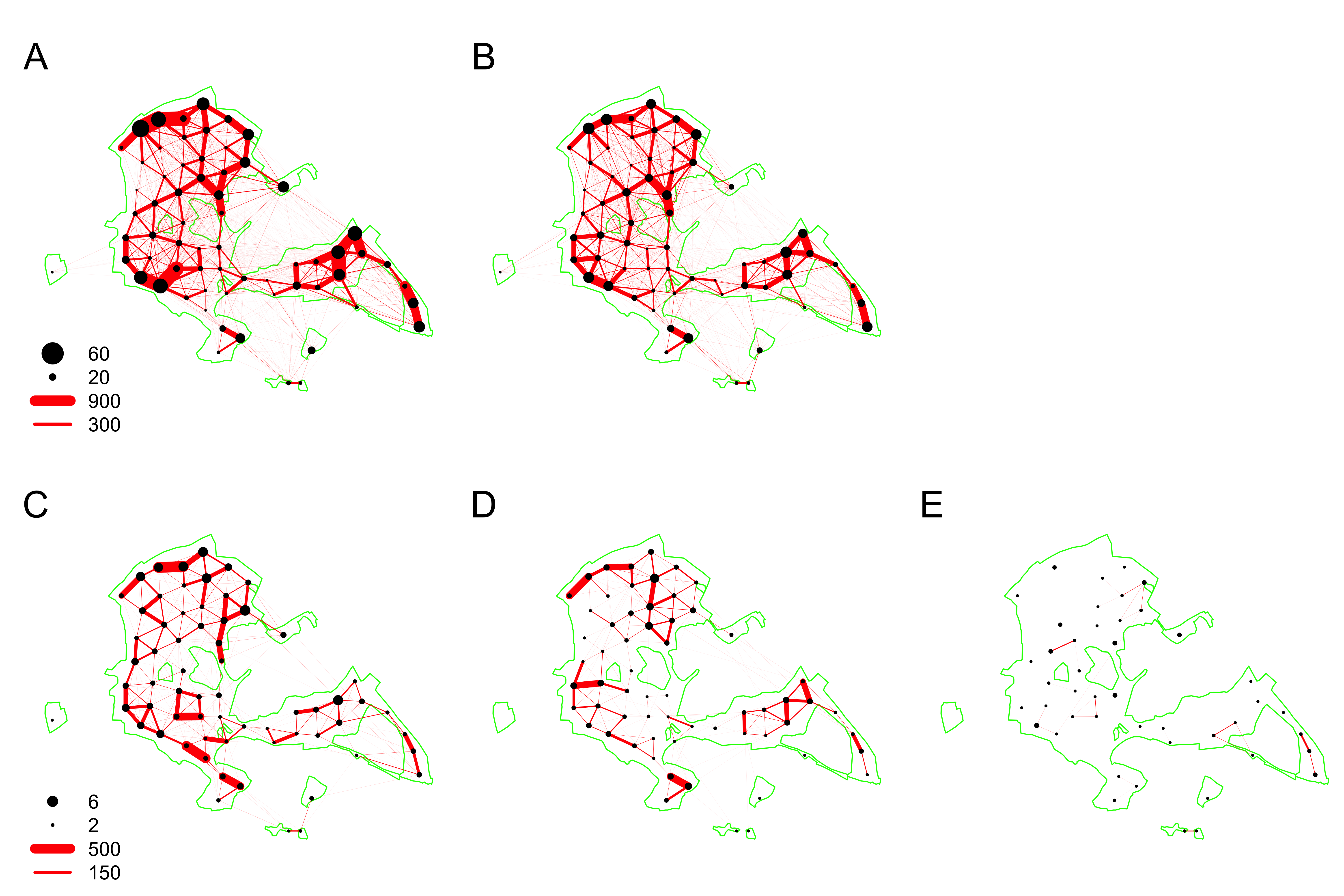

**Figure S2:** Movements broken down by species. (A) blue tits and (B) great tits show many more long-distance movements than (C) marsh tits, (D) coal tits, or (E) nuthatches. Red lines indicate observed daily movement between two feeders, and the thickness of the lines indicates the number of such detections. Black points represent feeder locations where individuals from that species were detected as having moved to or from, and the size of the black points indicates the number of individuals. There were approximately 10 to 20 times more individual great and blue tits (top row) than marsh tits, coal tits, and nuthatches, and thus the two rows are scaled differently. The green outline represents the outline of Wytham Woods and four small external woodplots

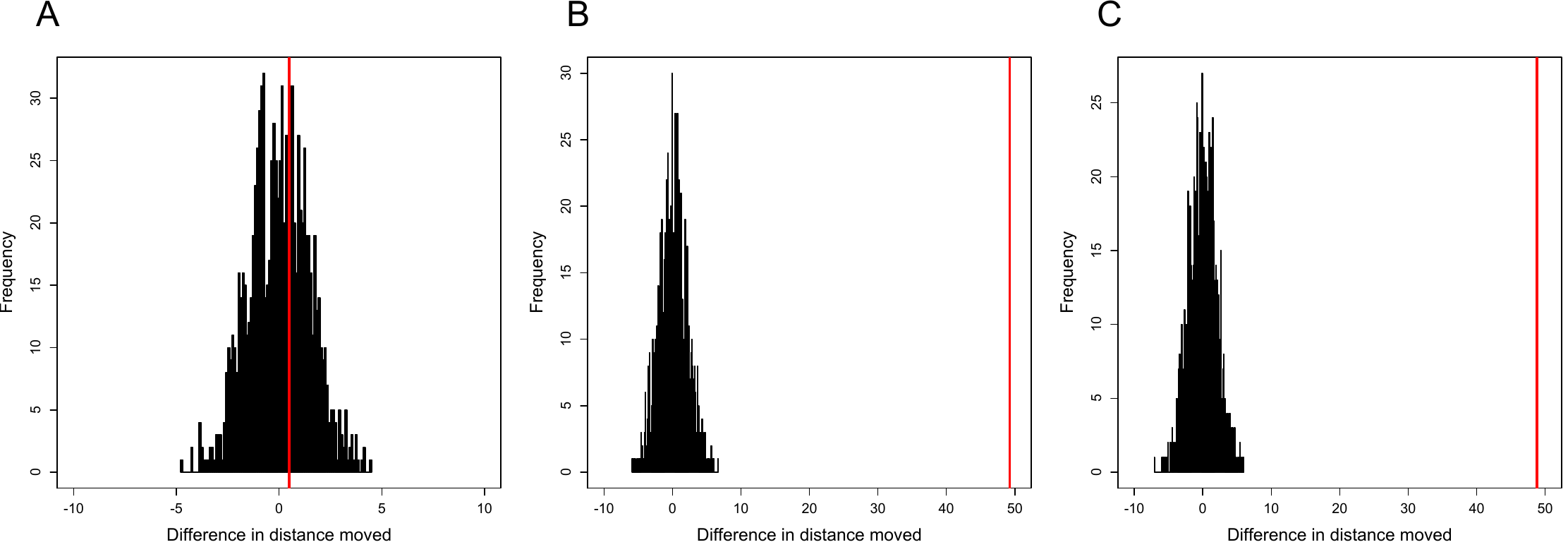

**Figure S3:** Analyses of the differences in the average distance moved between the three more common species: great tits, blue tits, and marsh tits. (A) Great and blue tits showed no difference in the average distance moved, but both made significantly longer movements on average than marsh tits. (B) shows blue versus marsh tits, (C) shows great versus marsh tits. Red line marks the observed difference and black histogram the expected difference from a null model shuffling the species identity of each record.

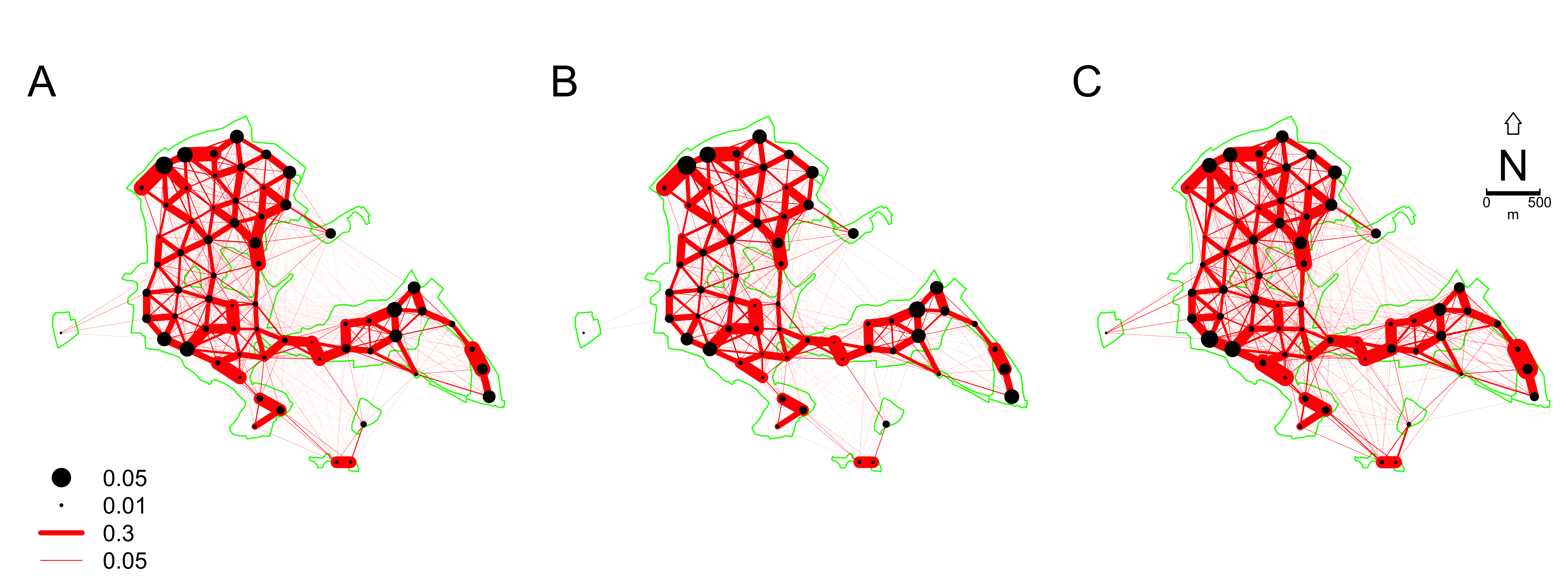

**Figure S4:** Probability of making a movement in a day between locations for (A) all birds, (B) adults, and (C) juveniles from all species over 4 winters of data. The thickness of each line represents the number of observations of a bird moving between the two feeding stations (black points) in the same day divided by the number of unique individuals seen at both sites combined. The size of the points represents the proportion of all unique individuals observed at each feeding station. The green outline represents the outline of Wytham Woods and four small external woodplots

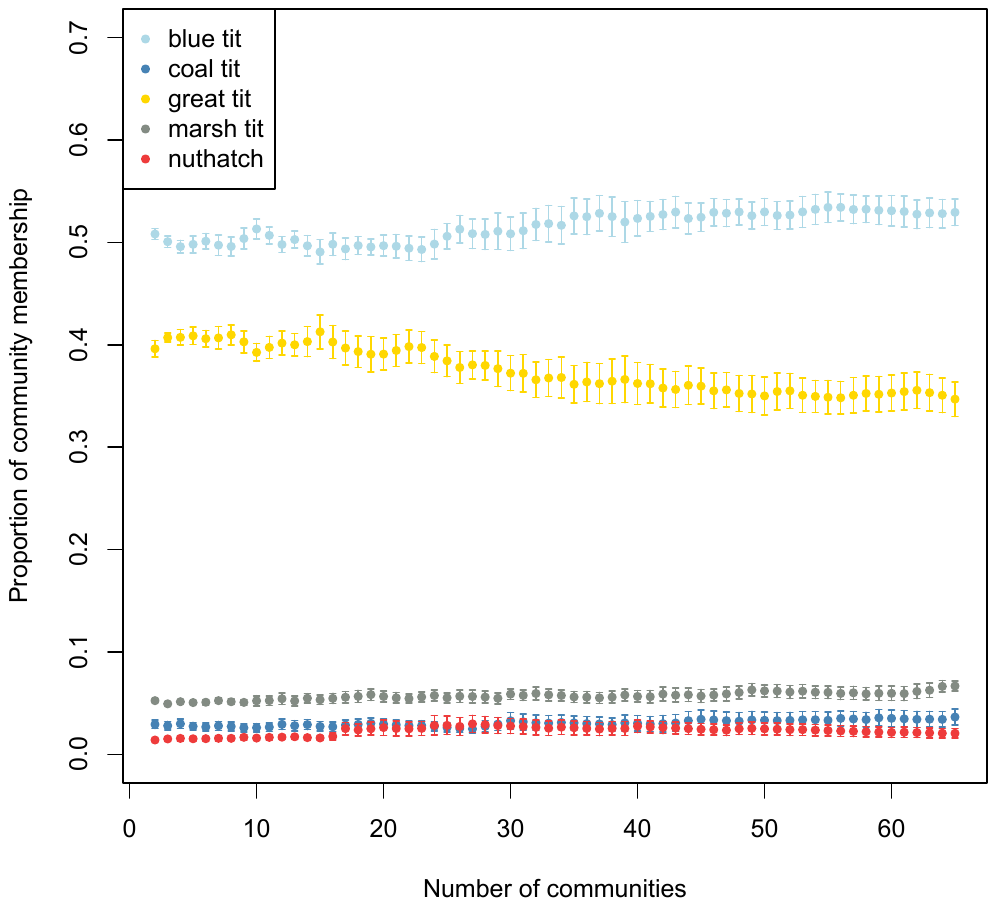

**Figure S5:** Average proportion of individuals of a species when the social network is partitioned into different communities shows that local community structure does not simply break down into species-level units as the proportion representation changes very little.
